# Camera trap based niche partitioning model of medium size ungulates and its implications on tiger’s prey availability in SE Asian rainforest

**DOI:** 10.1101/2021.08.20.457156

**Authors:** Andri Wibowo

## Abstract

The presence and survival of the Sumatran tiger were determined by the presence, density, and activity of its prey. Several medium size mammal belongs to the ungulate were known as the potential prey for the tiger. While, the information on the temporal availability of this prey in the rainforest is still limited. In here this study aimed to aims to model the niche partitioning of several mammalian species that was potential prey for tiger. The studied species were including barking deer (*Muntiacus muntjak*), bearded pig (*Sus barbatus*), and wild boar (*Sus scrofa*). The method was using camera trap with data analyses including the calculation of Kernel density, diel activity, and niche partitioning determined using overlap index. . Based on the results, barking deer and wild boar showed only singular activity peak while bearded pig has several activity peaks. Only wild boar showed a strict diurnal activity pattern between 06:00 h and 12:00 h. Barking deer showed a crepuscular behavior with several activities observed at 18:00 h. While bearded pig showed a nocturnal behavior and showed at least two peaks of activity, one between 09:00 to 13:00 and another between 18:00 to 24:00 h. Barking deer and bearded pig uses almost similar niches since those species have the highest overlap indices value equals 0.504(95%CI:0.193-0.824). The lowest overlap indices value was observed for barking deer and wild boar with overlap indice values of 0.032(95%CI:0.0-0.162). Considering the diurnal activity pattern of the tiger that is mostly active at day, then the available preys were either barking deer or bearded pigs. While since barking deer and bearded pigs were using the same niche, then there will be potential competition.

## Introduction

Understanding and assessing a species diel activity is fundamental to understanding the ecology and management of wildlife species. In this regard, activity curves (Werner & Anholt, 1993) can inform the movement ecology regulating the physiology of individuals and the growth of a population. This affects how a species makes decision and behaves following given tradeoffs related to forage acquisition, thermoregulation, and predator avoidance. Recently, an activity curve of multispecies representing relationships between species has been used to address hypotheses concerning interspecific and intraspecific competition and predator-prey interactions (Hernandez et al. 2015). Considering this is very important, a variety of methods have been proposed to estimate activity curves.

Previously, visual observations in the wild or in laboratory settings were used to assess and monitor the activities of an animal species. Whereas, visual observations in the wild contain obvious flaws related to concealment of the observer and detection of the subject. While, an observation in labs can result in altered species behaviors that preclude their widespread usefulness in determining activities of wild conspecifics. In regard to this situation, a new approach considered can be more effective and also accurate was using a camera trap.

Camera traps have more potential advantage to inexpensively and noninvasively monitor and assess the diel activity of wildlife populations. The equipment is relatively cheap and the technique does not rely on animal capture, which results in lower cost and labor effort, and minimize an adverse effect on species. Beside that camera trapping method may provide a means of gathering large datasets on elusive, rare, and protected species that can be used for a wide range of purposes. Camera traps have been used as a versatile research tool with wide ranges of utilities ranging from discovering new species, conducting simple vertebrate inventories, estimating density, measuring population dynamics, to examining habitat associations of entire animal communities (Kelly & Holub, 2008; Bridges & Noss, 2011) .

Sumatra rainforest is a suitable habitat for an endemic Sumatran Tiger. Existence and survival of this carnivore were related to the prey availability that mostly were mammals either small or medium sizes. Most preferred preys were species belong to ungulate groups. Among ungulate, tiger preys on wild boar (Dinata & Sugardjito, 2008). Despite tiger ecology and relationship with its prey have been widely documented and studied, an information on how the ecology of tiger prey is still limited. Here, this study aims to model the niche partitioning of several mammalia species that was potential prey for tiger.

## Methodology

### Study area

The study area was Bukit Tiga Puluh rainforest NP sizing 143,223 hectares located in eastern Sumatra. This park is consisting primarily of tropical lowland forest divided into a large portion in Riau Province and a smaller part of 33,000 ha in Jambi Province. This park is the last refuge of endangered species such as the Sumatran orangutan, Sumatran tiger, Sumatran elephant, and Asian tapir, as well as many endangered bird species.

### Camera trap assembly

The presence of medium size ungulates in study area were recorded by using camera traps (Figure 1) (Cheyne, 2012; Kuncahyo, 2017; Lashley et al. 2018; Subagyo, 2019). The camera traps were installed at tree trunks at 30-40 cm height. To allow the identification of the frontal and lateral body of species, the cameras were also positioned perpendicular to animal paths at 3-4 m distance. Likewise, the distance between one camera unit to another unit in grid cells was 1 km. In this study, the studied animals were including barking deer (*Muntiacus muntjak*), bearded pig (*Sus barbatus*), and wild boar (*Sus scrofa*).

**Figure 1.**
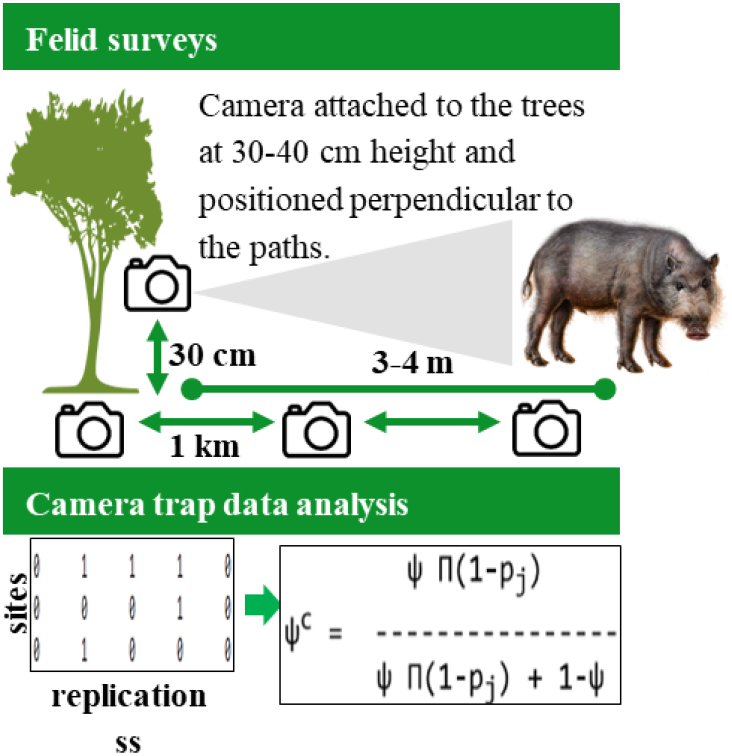
Camera trap assembly

### Kernel density and diel activity

Diel activity patterns were estimated using the kernel density method, which is ideal for circular data (Schmidt & Schmidt 2006; Ridout & Linkie 2009). For each ungulate species, the overall activity patterns were estimated, that is, pooling the data for each species. The Kernell density (Weglarczyk, 2018) was calculated as follow steps:

- let the series (x_1_, x_2_,…, x_n_) be an independent and identically distributed (iid) sample of n observations taken from camera trap x with an unknown probability distribution function f(x). Kernel estimate f(x) of original f(x) assigns each i-th sample data point x_i_ a function K(x_i_,t) called a kernel function in the following way:

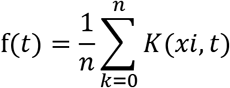
- with K(x_i_,t) is nonnegative and bounded for all x and t with 0 ≤ K (x_i_,t) < for all real x_i_,t and for all real x_i_.∫ K (x_i_,t)dt = 1
- Kernel transforms the point location of x_i_ into an interval centered (symmetrically or not) around x_i_.

### Niche partitioning

Ungulate species niche partitioning was determined using overlap coefficient (Santiago & Salvador 2017) that was defined as the proportion of the area under the curve that is superimposed between the two diel activity patterns. With Δ= 0 indicates no overlap indicating strictly nocturnal organism compared with a strictly diurnal one and Δ= 1 corresponds to a 100% overlap indicating daily activity patterns are identical (Meredith & Ridout 2016). The Δ confidence intervals were estimated with a bootstrap method, using 1000 random resampling of data with replacement (Meredith & Ridout 2014). The coefficient of overlapping ∆ for two probability density functions f(x) and g(x) is given by:

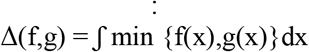

## Results and Discussions

The overall diel activity patterns and Kernel density for each of the ungulate species recorded in the Bukit Tiga Puluh Rainforest sites are available in Figure 2, 3, and 4. Barking deer and wild boar showed only singular activity peak while bearded pig has several activity peaks. Only wild boar showed a strict diurnal activity pattern between 06:00 h and 12:00 h. Barking deer showed a crepuscular behavior with several activities observed at 18:00 h. While bearded pig showed a nocturnal behavior and showed at least two peaks of activity, one between 09:00 to 13:00 and another between 18:00 to 24:00 h.

**Figure 2.**
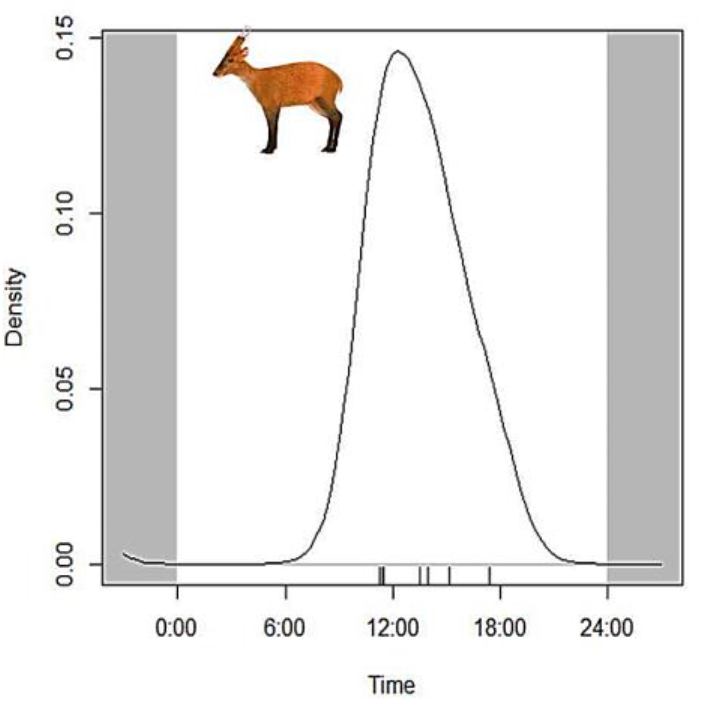
Kernel density and activity patterns of *Muntiacus muntjak* in the Sumatra rainforest. The strip at the base of the graphs represents the distribution of records for *M. muntjak*.

**Figure 3.**
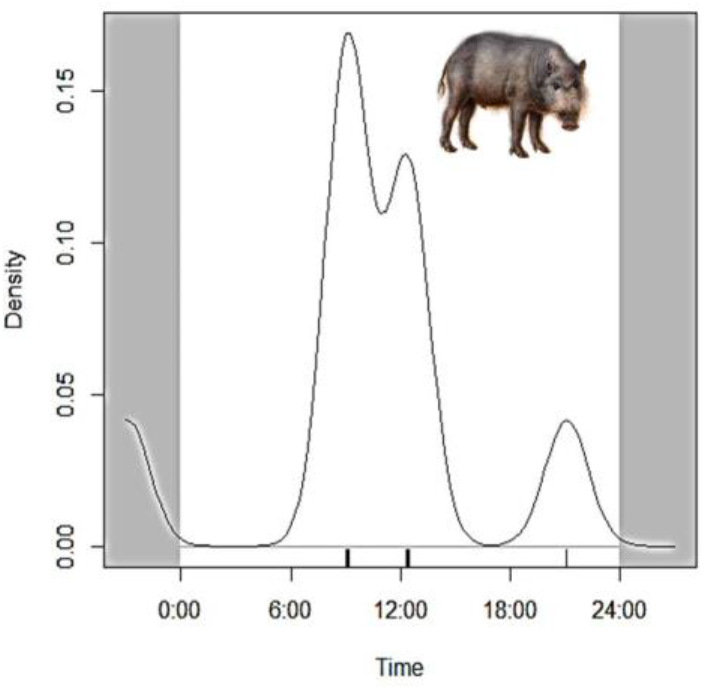
Kernel density and activity patterns of *Sus barbatus* in the Sumatra rainforest. The strip at the base of the graphs represents the distribution of records for *S.barbatus*

**Figure 4.**
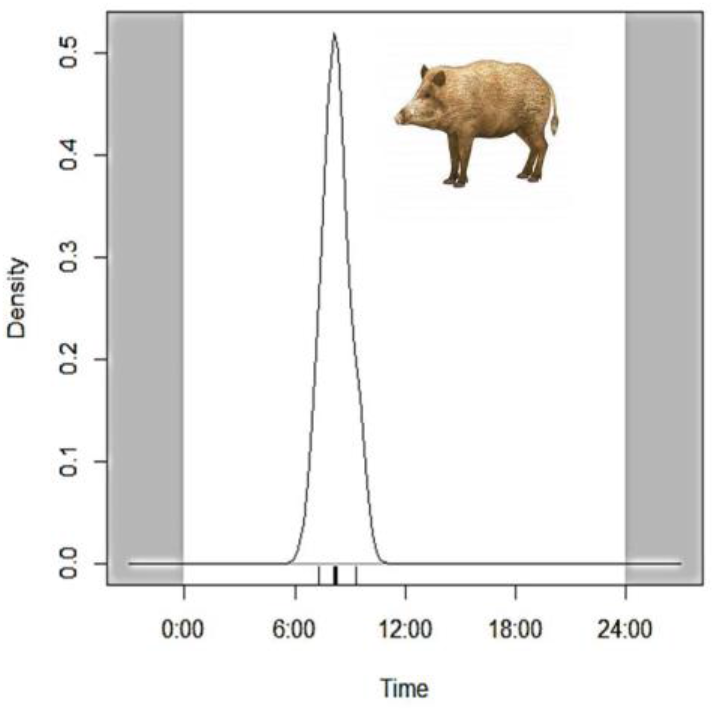
Kernel density and activity patterns of *Sus scrofa* in the Sumatra rainforest. The strip at the base of the graphs represents the distribution of records for *S. scrofa*.

There was evidence of correlative relationships between studied ungulates. The most significant correlations were observed for barking deer and bearded pig species (Figure 5). Those ungulates showed activity patterns with 50% overlap (95%CI:19–82%). Barking deer and bearded pig will use the same resources between 06:00 h and 18:00 h and also has the same activity peak at 12:00 h. In contrast, barking deer was not overlapped (Figure 6) with wild boar since the overlap indices value was very small equals only 3% (95%CI:0–16%). This indicates that barking deer has potential niche competition only with bearded pig. The second niche competitions (Δ = 0.426, 95%CI = 0.215-0.833) were observed between bearded pig and wild boar species (Figure 7). Those species were competed for same resources from 06:00 h to 12:00 h.

**Figure 5.**
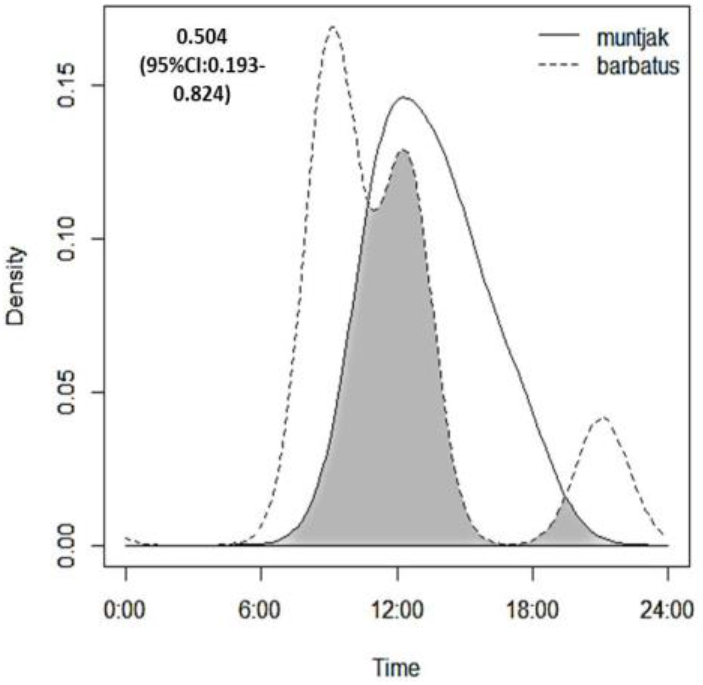
Niche overlap between *Muntiacus muntjak* and *Sus barbatus* in the Sumatra rainforest

**Figure 6.**
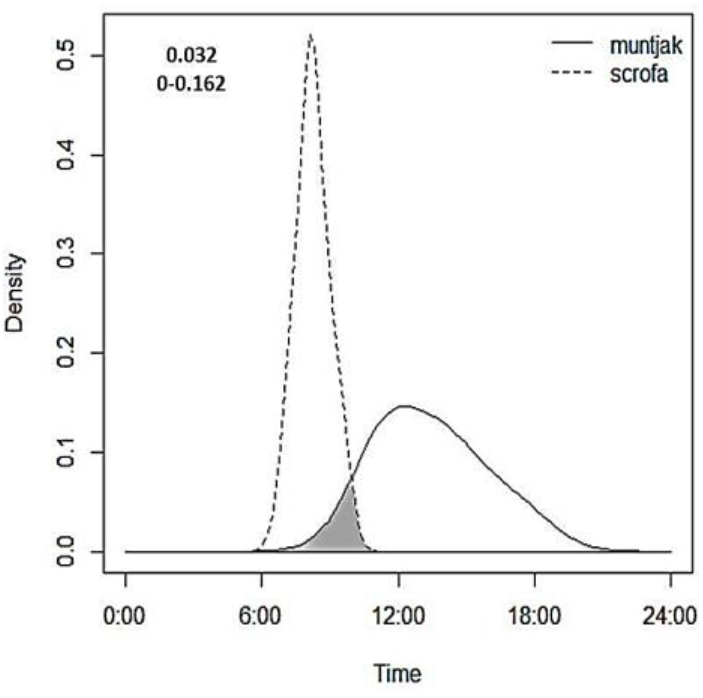
Niche overlap between *Muntiacus muntjak* and *Sus scrofa* in the Sumatra rainforest

**Figure 7.**
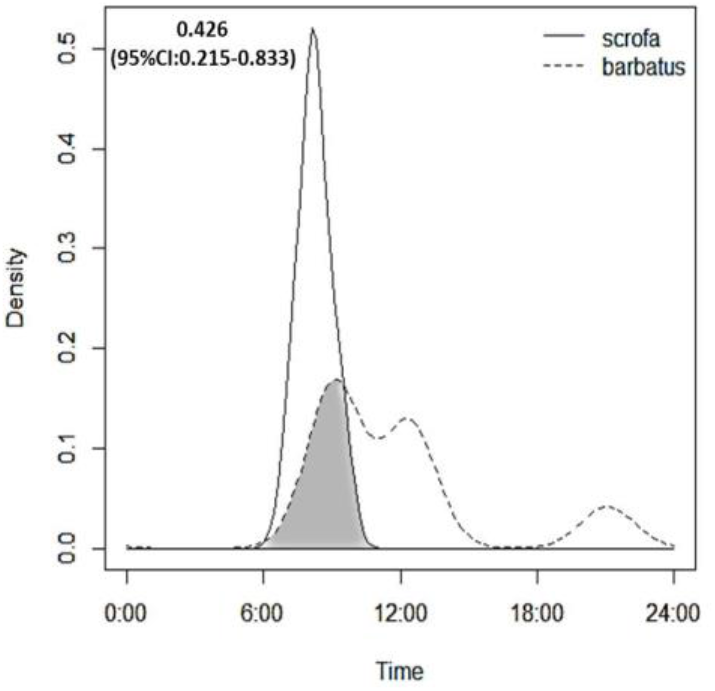
Niche overlap between *Sus barbatus* and *Sus scrofa* in the Sumatra rainforest

This study describes the first investigation into the activity patterns of a range of ungulate species on the tropical rainforest and adding to information gathered from across the Sumatra tiger species’ ranges and to those for which little is recorded. The study also contributes to the current literature that uses remote camera traps to infer activity and analyzed using overlap indices (Caravaggi et al. 2018). Findings in this result show a potential competition between barking deer and bearded pigs. This finding is consistent with other studies. Members of Suidae including bearded pigs are capable of competing with small to large-sized ungulates and feed on small vertebrates and invertebrates (Taylor & Hellgren 1997; Melberg 2012).

Those studied ungulate species were potential prey for the Sumatran tiger. Whereas, considering tiger activity that mostly were active at day from 06:00 to 18:00 h (Max et al. 2021), then only several ungulates were available as prey. Those ungulates including barking deer and bearded pig species. While wild boar cannot be potential prey since its activity pattern was not correlated with the tiger diel activity pattern.

## Conclusion

A competition involving ungulates was detected in the Sumatra rainforest. Barking deer and bearded pigs were the potential prey for tigers since those species were having overlapped niches with the tiger.

## References

Bridges, A. S. & Noss, A. J. 2011.Behavior and activity patterns. Camera-traps in Animal Ecology (eds A.F. O’Connell, J.D. Nichols & K.U. Karanth), pp. 57–70. Springer, New York

Caravaggi, A., Gatta, M., Vallely, M.C, Hogg, K., Freeman, M., Fadaei, E., Dick, J.T.A., Montgomery, W.I., Reid, N., Tosh, D.G. 2018. Seasonal and predator-prey effects on circadian activity of free-ranging mammals revealed by camera traps. PeerJ 6:e582.

Cheyne S, Ripoll B, Macdonald E, Sastramidjaja WJ. 2012. Standard operating procedure (sop) to install camera trap.

Dinata, Y. Sugardjito, J. 2008. The existence of Sumatran tiger (*Panthera tigris sumatrae* Pocock, 1929) and their prey in different forest habitat types in Kerinci Seblat National Park, Sumatra. Biodiversitas 9(3).

Downes, S. 2001.Trading heat and food for safety: costs of predator avoidance in a lizard. Ecol. 82: 2870–2881.

Santiago, E., Salvador, J. 2017. Hunters landscape accessibility and daily activity of ungulates in Yasuní Biosphere Reserve, Ecuador. Therya 8(1): 45–52.

Hernandez, F., Galvez, N. & Gimona, A. 2015. Activity patterns by two colour morphs of the vulnerable guiña, *Leopardus guigna* (Molina 1782), in temperate forests of southern Chile. Gayana 79: 102–105.

Kelly, M. J. & Holub, E. L. 2008.Camera trapping of carnivores: trap success among camera types and across species, and habitat selection by species, on Salt Pond Mountain, Giles County, Virginia. Northeastern Nat. 15: 249–262.

Kuncahyo. BA, Alikodra, HA, Gunawan, H. 2017. identifikasi faktor sebaran macan dahan (*neofelis diardi* cuvier, 1823) di ekosistem rawa gambut, taman nasional Sebangau.

Lashley, M.A., Cove, M.V., Chitwood, M.C. 2018. Estimating wildlife activity curves: comparison of methods and sample size. Sci Rep 8: 4173.

Max, A., Sibarani, M., Krofel, M. 2021. Predicting preferred prey of Sumatran tigers (*Panthera tigris sumatrae*) via spatiotemporal overlap. Oryx. 55. 197–203. 10.1017/S0030605319000577.

Melberg, S. 2012. Spatiotemporal competition patterns of Swedish roe deer and wild boar during the fawning season. Swedish University of Agricultural Sciences

Ridout, M. S., Linkie, M. 2009. Estimating overlap of daily activity patterns from camera trap data. Journal of Agricultural, Biological and Environmental Statistics 14:322–337.

Schmidt, F., Schmidt, A.. 2006. Nonparametric estimation of the coeficient of overlapping - Theory and empirical application. Computational Statistics and Data Analysis 50:1583–1596.

Subagyo, A. Muhammad Y., Sumianto Supriatna, J., Andayani, N., Mardiastuti, A., Sjahfirdi, L., Yasman, Sunarto. 2019. Survei dan monitoring kucing liar (carnivora:felidae) di taman nasional Way Kambas, Lampung, Indonesia.

Taylor, R. B. Hellgren, E. C. 1997. Diet of feral hogs in the western south Texas plains. South-western Naturalist 42: 33–39.

Weglarczyk, S. 2018. Kernel density estimation and its application. ITM Web of Conferences. 23. 00037.

Werner, E. E. & Anholt, B. R. 1993. Ecological consequences of the trade-off between growth and mortality rates mediated by foraging activity. Am. Nat. 142: 242–272.

